# Expanding large-scale mechanistic models with machine learned associations and big datasets

**DOI:** 10.1101/2022.11.21.517431

**Authors:** Cemal Erdem, Marc R. Birtwistle

## Abstract

Computational models that can explain and predict complex sub-cellular, cellular, and tissue level drug response mechanisms could speed drug discovery and prioritize patient-specific treatments (i.e., precision medicine). Some models are mechanistic: detailed equations describing known (or supposed) physicochemical processes, while some models are statistical/machine learning-based: descriptive correlations that explain datasets but have no mechanistic or causal guarantees. These two types of modeling are rarely combined, missing the opportunity to explore possibly causal but data-driven new knowledge while explaining what is already known. Here, we explore a combination of machine learning with mechanistic modeling methods to develop computational models that could more fully represent cell-line-specific drug responses. In this proposed framework, machine learning/statistical models built using omics datasets provide high confidence predictions for new interactions between genes and proteins where there is physicochemical uncertainty. These possibly new interactions are used as new connections (edges) in a large-scale mechanistic model (called SPARCED) to better recapitulate the recently released NIH LINCS Consortium large-scale MCF10A dataset. As a test case, we focused on incorporating novel IFNγ/PD-L1 related associations into the SPARCED model to enable description of the cellular response to checkpoint inhibitor immunotherapies. This work is a template for combining big data, machine-learning-inferred interactions with mechanistic models, which could be more broadly applicable towards building multi-scale precision medicine and whole cell models.

## Introduction

The molecular signaling mechanisms of cancer cells are highly heterogenous, leading to treatment adaptations and recurrence. Thus, the need for personalized interventions to block tumor growth has emerged. The traditional drug discovery pipeline is comprised of extensive trial-and-error experiments, testing thousands of chemicals, refining their structure for safety and toxicity, and administering years of clinical trials. This burden can be reduced by understanding the underlying molecular mechanisms with the help of computational models (1,2).

Computational tools and models are becoming indispensable in medical research, where a cycle of experimentation and computation is used to learn about and test new hypotheses. The models guide experimental hypothesis generation, and experimental observations enable fine-tuning computational models to understand the biological phenomena. Owing to the advances in wet-lab experimental techniques and tools, “Big Data” repositories become more prominent each year. The knowledge base of these databases includes genomics, proteomics, epigenetics, and clinical information (3–8). To understand the underlying biological facts, analysis of the wealth of the aforementioned big datasets should become more practical and go beyond context-dependent and scope-limited biological events.

Building computational models that explain and predict such highly heterogenous and complex cellular responses is no easy task. The popular mechanistic models are sets of detailed equations describing curated knowledge of what is happening within the cells. Such models (9–11) are usually small in scale: tens of equations and 10s-100s of model species (Fig. 1) Another popular class is machine learning based models, which are data-driven, descriptive, and mostly large-scale (genome-wide or exome-wide) (2,12–14). These types of models are generally coined as black-box models because although they perform well in precision/recall metrics, how they do so is blurry (Fig. 1). So far in the literature, these two types of models are rarely combined, missing the opportunity to generate new knowledge while explaining what is already known (15).

**Fig. 1.**
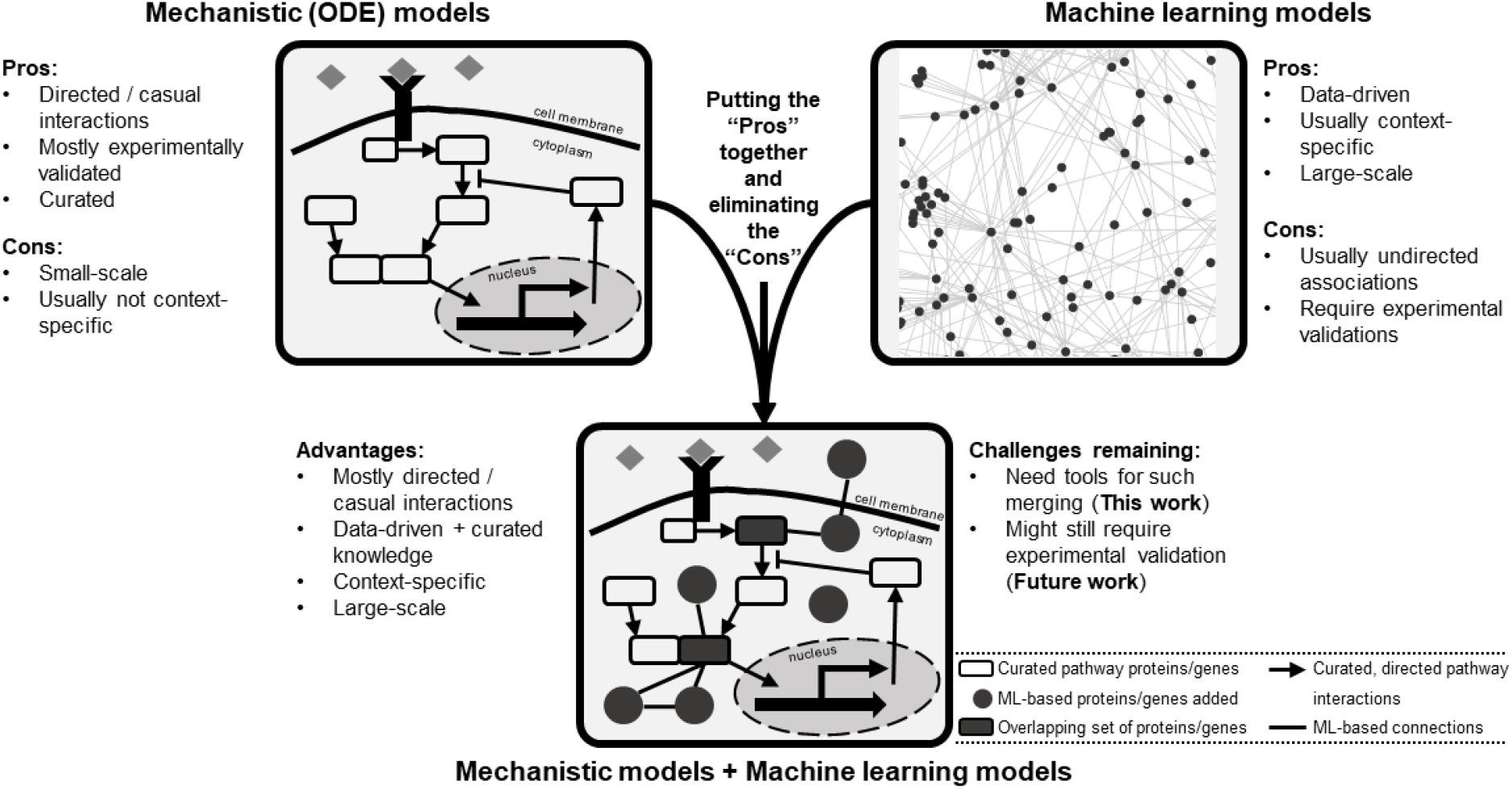
Different computational modeling types of biological data possess a variety of pros and cons and provide an opportunity for model merging. The mechanistic models are mostly curated, usually small-scale, causal networks of signaling pathways. Machine learning models are data-driven, large-scale, and usually correlative associations. Combining these two modes of modeling provides an opportunity for creating larger scale data-informed models to generate novel hypotheses for experimental validation. The merged model would include curated lists of pathway genes (species) as well as genes of new connections inferred via machine learning models. The final model structure could represent a collection of overlapping genes (and gene products) and interactions present in both lists.

Here, we explore a combination of both methods to develop better models that will accurately represent generated biological knowledge. Connections inferred via a machine-learning pipeline (called MOBILE (16)) are inserted as new interactions into a large-scale mechanistic model (called SPARCED (17)) to better recapitulate the recently released MCF10A dataset (18). The NIH-LINCS Consortium and MCF10A Common Project recently released this dataset, consisting of multiple omics assay types on breast epithelial MCF10A cell line. MOBILE is a new pipeline to integrate multi-omics datasets and identify context-specific interactions. SPARCED is one of the largest mechanistic models of mammalian cells and is an open-source, human-interpretable, and easy to alter modeling format. Previously, we used MOBILE to obtain candidate context-specific associations in MCF10A cells and proposed novel sub-networks of IFNγ / PD-L1 regulation and differential pathway activities under TGFβ1 and BMP2 stimulation. Here we focused on incorporating novel IFNγ/PD-L1 related associations into the SPARCED model to enable description of the cellular response to checkpoint inhibitor immunotherapies. Using this work as a recipe for combining big data, machine-learning-inferred interactions with mechanistic models, we can start building multi-scale precision medicine and whole cell models.

## Materials and methods

In this work, we use ligand-specific interactions between genes as new connections in a large-scale mechanistic model to study the effect of the newly added gene interactions in model responses.

### MOBILE

It is a recent tool for finding context-specific network features by integrating pairs of omics datasets. In short, statistical associations are calculated between pairs of chromatin accessibility regions, mRNA expressions, and protein/phosphoprotein levels. Lasso (least absolute shrinkage and selection operator) regression models are run in replicate to select coefficients with high occurrence rates (16,19,20). The so-called Integrated Association Networks (IANs) are generated by combining the association networks inferred for RPPA+RNAseq and RNAseq+ATACseq data inputs. Finally, the IANs are coalesced into gene-level networks: nodes representing genes of the assay analytes and edges representing the inferred Lasso coefficients.

### SPARCED

The starting mechanistic model used in this work is obtained from the SPARCED repository (github.com/birtwistlelab/SPARCED/tree/develop) (17). It is a recent framework for large-scale mechanistic modeling that enables model file creation using simple text files as input with minimal coding requirements. In short, a set of annotated text files are constructed to define model specifics. Then, Jupyter notebooks are used to process these files and create community-standard model file type called SBML (21,22). The software was first built to replicate the largest mammalian single-cell mechanistic model of proliferation and death signaling (9,17). Then, an expanded SPARCED model was created to include IFNγ signaling sub-module and the new model was named as SPARCED-I-SOCS1 (17). This final model and its input files are used as the basic model in this work and is modified further with the MOBILE inferred set of new connections.

### MEMMAL

#### (1) MEMMAL enlargeModel notebook

The Jupyter notebook processes the machine learning model inferred connections list and creates necessary Species (genes, mRNAs, proteins, phosphoproteins), RateLaws (the reaction format and related parameters), Gene Regulatory Interactions (defining transcriptional activators and repressors) and finds relevant new omics data from LINCS datasets. The input files for SPARCED pipeline are then updated with new rows and columns followed by default model compilation and simulation steps.

#### (2) MEMMAL createModel_o4a notebook

The recently released Jupyter notebook to create an integrated SBML version of the SPARCED type models. Creating the model file fully in SBML format provides extensive speed-up of simulations. The newly updated input files are used to create and compile the MEMMAL model.

#### (3) MEMMAL createModel notebook

The default SPARCED notebook to create hybrid model format. Although not used in this work, users can re-compile the model and run hybrid simulations to explore single-cell responses of IFNγ / PD-L1 model.

#### (4) MEMMAL fitModel notebook

The Jupyter notebook provides updated parameter values and code to run model verification simulations.

#### (5) MEMMAL runModel notebook

The Jupyter notebook is used to simulate and explore multiple scenarios for the new MEMMAL model.

### Code availability

MEMMAL code is available at the GitHub repository github.com/cerdem12/MEMMAL.

### Data availability

All the data used in this study are available within the MOBILE repository and adapted from (18).

## Results

### Large-scale mechanistic models can become larger and more precise by expansion using machine learned relationships

There are only a handful of large-scale (hundreds of genes, thousands of species) mechanistic signaling pathway models in the literature (10). Usually, such big models are constructed by bottom-up modeling or by semi-manual stitching of previously published models (9). Both approaches are time consuming, manually curated, and biased for including/excluding model components: genes, proteins, post-translational modifications, interactions, or even cellular compartments. Here, we tackle this “what-to-add” problem by using association networks inferred via data-driven machine learning algorithms. Specifically, we used two recent tools for network inference and mechanistic model creation.

The Mechanistic Modeling with Machine Learning (MEMMAL) tool presented here (Fig. 2) is comprised of scripts to expand mechanistic models created using SPARCED pipeline (17) with candidate connections generated by the tool called MOBILE, a recent pipeline for multi-omics data integration (16). The MEMMAL software is based on the SPARCED GitHub repository (github.com/birtwistlelab/SPARCED/tree/develop), which enables users to easily create, alter, and simulate mechanistic large-scale models. Here, we combine novel connections inferred via MOBILE with a large-scale mechanistic model using SPARCED. Specifically, we add an immune-checkpoint related sub-module to the existing pan-cancer model to study effects of the newly added gene products on the regulation of Interferon Regulatory Factor 1 (gene name IRF) and Programmed Death Ligand 1 (PD-L1, gene name CD274) upon interferon-gamma (IFNγ) stimulation.

**Fig. 2.**
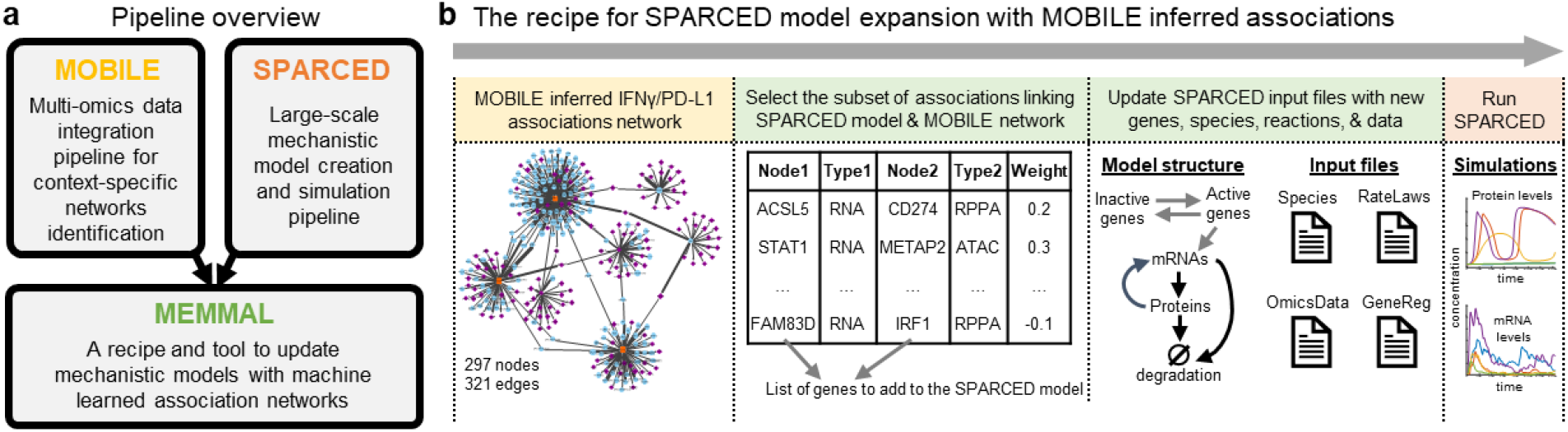
MEMMAL is a pipeline to merge mechanistic modeling with machine learning. **a** The MEMMAL pipeline combines mechanistic models created by SPARCED with association networks generated via MOBILE pipeline. **b** The recipe for MEMMAL pipeline starts by obtaining a set of connections not presented in the candidate mechanistic model. Here, the novel gene-level connections list is inferred via the MOBILE tool and then filtered for overlap with SPARCED model genes. Next, this candidate network is imported into SPARCED environment, where the MEMMAL `enlargeModel` Jupyter notebook processes the network file and updates SPARCED input files The nodes (genes) of the IFNG / PD-L1 subnetwork are used to create new genes and species (mRNAs, proteins, phosphoproteins) for SPARCED. The new genes can get activated/inactivated as described in SPARCED. The expanded MEMMAL model is created and compiled by default SPARCED model notebooks. The final step in MEMMAL is to run user defined exploratory simulations to gain insights on the effects of new connections added.

### MOBILE pipeline integrated LINCS MCF10A multi-omics dataset to infer ligand-specific associations

The normal-like breast epithelial cell line MCF10A was recently profiled with multiple assay types under multiple ligand stimulation conditions (18). Using this newly released multi-omics dataset, our lab introduced the MOBILE pipeline for data integration and showed how ligand-specific associations can be inferred (16). One of the ligands included in the LINCS study that induced MCF10A growth inhibition was interferon gamma (gene name: IFNG), where Gross et al. showed an overall strong signal with 24- and 48-hour stimulation. So, we previously analyzed the LINCS MCF10A dataset to find IFNγ-specific associations to find novel connections with PD-L1 axis (16). IFNγ can induce transient PD-L1 (gene name CD274) expression, a transmembrane protein that binds to its receptor PD-1 on T-cells (23–25). This binding inhibits tumor clearance, where targeted therapies towards these proteins are a new class of anti-cancer drugs: the immune checkpoint inhibitors (26). However, inter- and intra-tumor variability of PD-L1 expression results in heterogeneous patient responses and makes the response predictions a challenge (27). A more thorough understanding of the regulatory mechanism of PD-L1 expression would then provide insights to identify new immunotherapeutic drugs or treatment options.

Applying MOBILE, we generated data-driven IFNγ-specific integrated associations network, which had 297 nodes (genes) and 321 edges (connections) (Fig. 2b and Supplementary Fig. 1). We further filtered this network by looking for connections with STAT1 (the only overlapping gene with the mechanistic model). The final list of candidate connections had nine genes (ACSL5, BST2, CD274, CLIC2, FAM83D, HIST2H2AA3, IRF1, METAP2, and STAT1) and 14 connections. The list is imported into the SPARCED environment to start altering the existing mechanistic model structure (Fig. 2b and Supplementary Fig. 1).

### SPARCED modeling makes mechanistic model expansions easy

SPARCED is a recent software (17) and modeling framework for large-scale mechanistic modeling. It enables SBML model file creation using simple text files as input with minimal coding requirements. Jupyter notebooks (28) are used to process the input files and to create the model files. The software was first built to replicate the largest mammalian single-cell mechanistic model of proliferation and death signaling (9) and was then expanded to include a new sub-module of IFNγ signaling (29). So, the starting mechanistic model in this work, SPARCED-IFNG-SOCS1 already includes IFNγ submodule (Fig. 3a, gray background), with a total of 149 genes, 1302 species, and 3584 ratelaws.

**Fig. 3.**
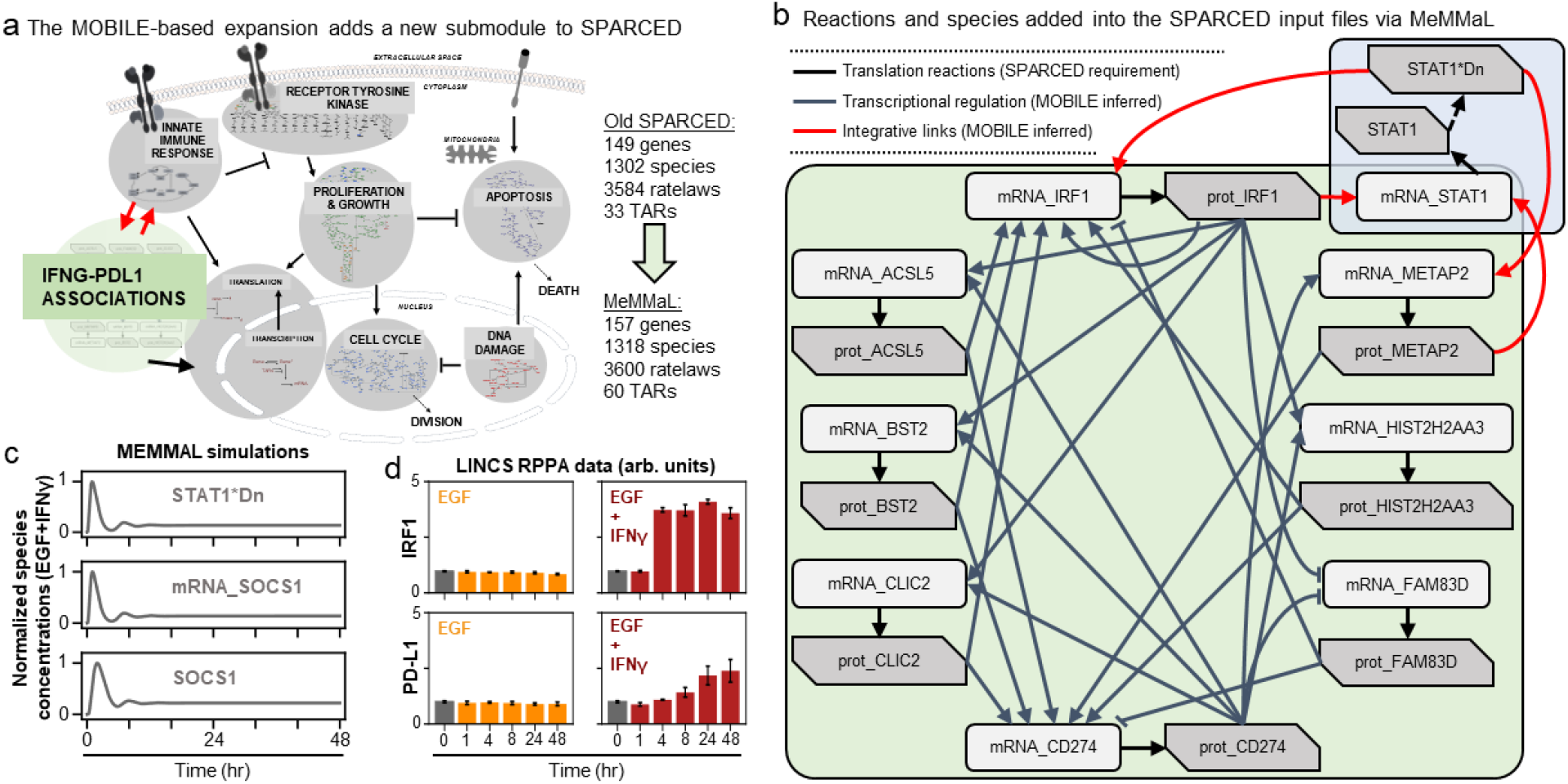
MOBILE inferred IFNG / PD-L1 network nodes and connections are inserted into the SPARCED-IFNG model using MEMMAL. **a** The reactions added into the SPARCED include translation (black arrows). The connections from MOBILE are translated as transcriptional activation and repression (TAR) reactions in MEMMAL (gray arrows). The TAR reactions linking existing SPARCED species with the newly added species are represented as integrative links (red arrows). **b** The SPARCED network is enlarged to include a sub-network spanning innate immune response and PD-L1 regulation. The final MEMMAL model is 157 genes, 1318 species, and 3600 ratelaws, 60 TARs, and 3885 parameters large. **c** The final MEMMAL model shows canonical IFNγ response with expected STAT1 and SOCS1 dynamics. Normalized simulation trajectories of the activated nuclear STAT1 dimer (STAT1*Dn), SOCS1 mRNA (mRNA_SOCS1), and free SOCS1 protein (SOCS1) are shown (solid gray lines). d IFNγ (maroon bars) induces IRF1 (top row) and PD-L1 (bottom row) expression as opposed to no induction by EGF-only stimulation (orange bars). The normalized protein levels from the LINCS dataset are shown. Error bars represent standard deviation of the triplicates.

### MEMMAL incorporates MOBILE-inferred gene-level statistical associations into SPARCED as gene regulatory mechanisms

The list of candidate connections from MOBILE pipeline are processed via MEMMAL `enlargeModel` notebook to add rows and update SPARCED input files (Fig. 2b). As a default SPARCED requirement, each gene node from MOBILE list is interpreted to create active gene, inactive gene, mRNA, and protein species, with relevant basic reactions: gene switching, transcription, translation (Fig. 3b, black arrows), mRNA degradation, and protein degradation. Importantly, the MOBILE inferred connections (14) are interpreted as transcriptional activator and repressor (TAR) reactions (Fig. 3b) because the MOBILE inferred connections are obtained by looking at pairs of mRNA-protein and chromatin region-mRNA dataset pairs. A logical way a protein affecting another mRNA’s expression level is by transcriptional regulation. Additionally, a highly open chromatin region can induce replication, which potentially yields higher mRNA expression and thus another gene regulatory connection. Thus, all the candidate associations are treated as TARs in the current MEMMAL pipeline. For future work, users should decide how to handle such connections.

The negative valued associations here are treated as inhibitory whereas the positive magnitude connections are added as activators (Fig. 3b, gray and red arrows). Some of the transcriptional activators are labeled as integrative links because they connect existing SPARCED model genes with the new gene species (Fig. 3b, red arrows). After all the input files are updated, `createModel_o4a` Jupyter notebook is used to create and compile the MEMMAL SBML model file (Fig. 2b). The MEMMAL expansion of SPARCED via MOBILE inferred network resulted in addition of eight genes, 16 species, 16 signaling reactions, and 27 transcriptional regulatory mechanisms (Fig. 3a). With the current addition, the SPARCED model now includes an IFNγ-PD-L1 submodule (Fig. 3a, green background).

After the model expansion, we first verified the model can recapitulate previous observations (Fig. 3c). We show that inclusion of new species and reactions did not alter canonical STAT1 / SOCS1 response to IFNγ stimulation. Then, the next step was to fit parameters for new reactions added to recapitulate the experimental data from LINCS RPPA assay (Fig. 3d). It was shown that both IRF1 and PD-L1 are induced only by IFNγ (+EGF) when compared to EGF-only stimulation. So, MEMMAL `fitModel` Jupyter notebook provides updated parameter values and scripts to compare simulation trajectories with LINCS datasets (Fig. 3c and 4a).

**Fig. 4.**
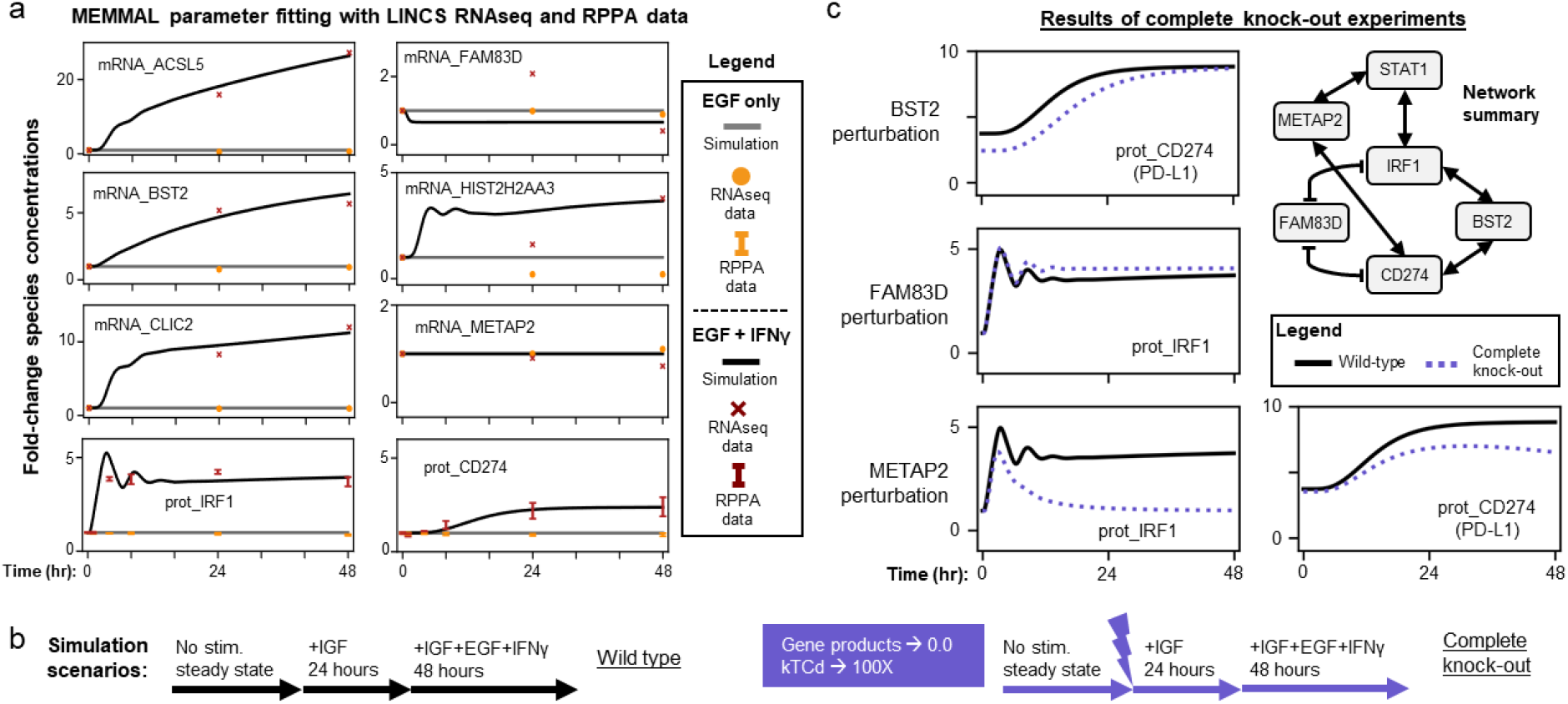
MEMMAL can replicate the previous SPARCED-IFNG model and offers new insights into IFNG regulation of IRF1 and PD-L1 dynamics. **a** MEMMAL model parameters are fitted to recapitulate experimental data from LINCS RNAseq and RPPA assays. Fold-changes are shown for data (dots, crosses, and error bars, STD) and simulations (solid lines). Most mRNAs and IRF1 and PD-L1 (gene name CD274) are induced by IFNγ. **b** Simulation scenarios to test the effects of newly added genes. The parameter fitted MEMMAL model is simulated with reported perturbation under IGF1 stimulation (basal growth condition) and then stimulated with EGF+IFNγ for 48 hours. **c** Comparison of complete gene knock-out perturbation scenario (dotted lines) to wild-type (no perturbation, black lines) condition shows genes with induced IRF1 and PD-L1 changes. Among the newly added genes, METAP2 induces the greatest change: a complete recession of IRF1 response and decreased PD-L1 steady-state level. The network diagram (summary of Fig. 3b) shows the connections among shown genes and STAT1.

### MEMMAL model offers exploration of the effect of novel connector genes on the expression of PD-L1 expression in response to IFNG

After we show the new MEMMAL model simulations closely follow experimental mRNA and protein levels (Fig. 4a), we simulated multiple scenarios (Fig. 4b) to explore the effects of the novel genes and gene products added (Fig. 4b) on the IRF1 and PD-L1 responses. We wanted to find new connections important in regulating PD-L1 expression to be able to start pinpointing mechanisms for better immunotherapies. By comparing “wild-type” simulations to “complete knock-out (protein, gene, and mRNA levels set to zero)” condition, we compiled the observed changes in IRF1 and PD-L1 protein levels (Figs. 4b-c).

Perturbing BST2 caused a small decrease in initial PD-L1 levels, which later reaches to wildtype response levels (Fig. 4c, top row). Similarly, perturbing FAM83D resulted in changes only in IRF1 levels (Fig. 4c, middle row). The IRF1 levels were slightly increased. The METAP2 perturbations caused a significant change in late IRF1 responses, with additional PD-L1 level decrease (Fig. 4c, bottom row). For all these scenarios, the levels of all other newly added mRNAs and proteins were unchanged compared to wild-type, and thus not shown here.

Note that we observed the largest changes induced after METAP2 perturbation, which has constant expression levels across 48 hours of stimulation (Fig. 4a). This is important because it provides evidence that mechanistic models can be utilized to prioritize experimental candidates by testing multiple in silico perturbations and merging mechanistic models with machine learning derived connections can help start learning new biology.

## Discussion

Combining and synergizing machine learning with mechanistic modeling would bring clinically predictive computational models and personalized medicine to life. To that end, here we introduced a recipe to expand a large-scale mechanistic model with machine learned connections between gene products. Because understanding PD-L1 regulation mechanisms would help us design better therapeutic interventions, we focused on exploring the IFNγ / PD-L1 axis. We used the LINCS MCF10A dataset and added the recently inferred (via MOBILE pipeline) IFNγ / PD-L1 connections to the existing SPARCED mechanistic model. We then were able to study the effects of new gene regulatory mechanisms. We showed that perturbing BST2, FAM83D, or METAP2 induces changes in PD-L1 and IRF1 dynamics.

MEMMAL could serve as an initial step towards combining mechanistic models with machine learning by providing a rationale for such a merging protocol. MEMMAL protocol first creates genes and gene products (mRNA and protein) if MOBILE list nodes are not present in SPARCED. It then updates -omics level information for the new genes and adds corresponding reactions. It also assigns transcriptional activator and repressors (based on MOBILE association coefficient signs) and related rate constant parameters. The updated SPARCED input files are then processed via modified default Jupyter notebooks to simulate desired simulations. Although MEMMAL makes use of recent tools from our lab, the idea is applicable to other tools available in the literature. For instance, rule-based modeling software like BioNetGen (30) and PySB (31) can also be used for mechanistic model creation and update if machine learning predicted associations are converted into new rules. Another possible application can include INDRA (32) if the new connections are put into suitable sentence format. Such options will be valuable to expand the MEMMAL idea and its applications.

The MOBILE pipeline was used to infer ligand-specific and statistically robust association networks (16). Here we used a filtered list of connections for interferon-gamma signaling and among them some genes were already shown to be associated with immunotherapeutic signatures including BST2, CLIC2, and FAM83D (33–37). In short, BST2 is part of an anti-CTLA4 response in melanoma (33) and CLIC2 is a favorable prognosis biomarker (34). FAM83D functions in cell growth regulation and is a prognostic marker for multiple cancer types (35,37). In addition to such pieces of literature support, we can take a step further explore their mechanistic functionalities by combining these genes and their predicted connections as new interactions in a computational model.

Here introduced MEMMAL provides a starting point for merging a recent large-scale mechanistic modeling tool with another recent multi-omics data integration pipeline. We then used MEMMAL to test novel candidate interactions for their effect on regulating IRF1 and PD-L1 expression and found that METAP2 is a good candidate yet to be studied experimentally. We believe combining big data, machine learning, and mechanistic models provides an invaluable asset to unravel novel context-specific mechanisms.

## Author Contributions

Conceptualization, C.E. and M.R.B.; Methodology, C.E. and M.R.B.; Software, C.E.; Validation: C.E.; Formal analysis: C.E.; Resources: M.R.B.; Writing – Original Draft: C.E. and M.R.B.; Writing – Review & Editing: C.E. and M.R.B.; Visualization: C.E. and M.R.B.; Supervision: C.E. and M.R.B.; Project administration: C.E. and M.R.B.; Funding acquisition: M.R.B.

## Acknowledgements

The authors acknowledge funding from the National Institutes of Health Grants 1R35GM141891 and U54HG008098-LINCS Center (M.R.B.). C.E. was an NIH-LINCS Consortium Postdoctoral Fellow (2018-2020).

