## Supplementary figures and images for "Expanding large-scale mechanistic models with machine learned associations and big datasets"

### Supplementary Figure 1

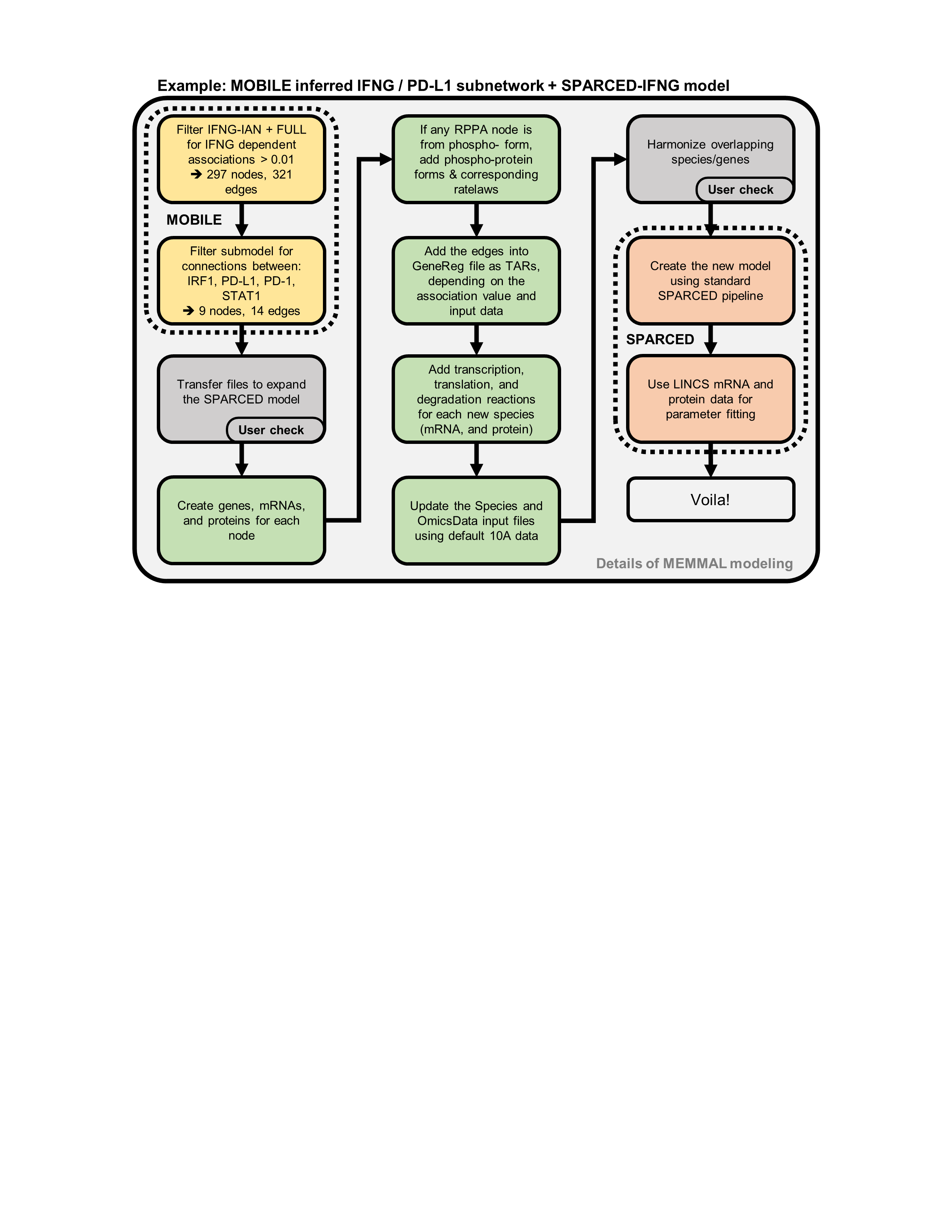
